# The genome of the blind bee louse fly reveals deep convergences with its social host and illuminates Drosophila origins

**DOI:** 10.1101/2022.11.08.515706

**Authors:** Héloïse Bastide, Hélène Legout, Noé Dogbo, David Ogereau, Carolina Prediger, Julie Carcaud, Jonathan Filée, Lionel Garnery, Clément Gilbert, Frédéric Marion-Poll, Fabrice Requier, Jean-Christophe Sandoz, Amir Yassin

## Abstract

The nests of social insects often harbor a rich fauna of intruders known as inquilines.^1^ Social inquilines are usually closely-related to their host due to potential genetic predispositions,^2,3^ but how phylogenetically distant non-social inquilines adapt to their hosts remains unclear. Here, we analyzed the genome of the wingless and blind bee louse fly *Braula coeca*, an inquiline kleptoparasite of the Western honey bee *Apis mellifera*.^4,5^ Using large phylogenomic data, we confirmed recent accounts that the bee louse fly is an aberrant drosophilid,^6,7^ and showed that it had likely evolved from a sap-breeder ancestor associated with honeydew and scale insects wax. Unlike many parasites, such as the human louse, the bee louse fly genome did not show significant erosion or strict reliance on an endosymbiont, likely due to a relatively recent age of inquilinism. However, a striking parallel evolution in a set of gene families was observed between the honey bee and the bee louse fly. Convergences included genes potentially involved in metabolism and immunity and the loss of nearly all bitter-tasting gustatory receptors in agreement with life in a protective nest and a major diet of honey, pollen, and beeswax. Vision-related and odorant receptor genes also exhibited rapid losses. Only genes whose orthologs in the closely related *Drosophila melanogaster* respond to honey bee pheromones components or floral aroma were retained, whereas the losses included orthologous receptors responsive to the anti-ovarian honey bee queen pheromones. These results indicate that deep genomic convergences can underlie major morphological and neuroethological transitions during the evolution of inquilinism between non-social parasites and their social hosts.

## Results and Discussion

### The bee louse fly Braula coeca is an aberrant member of the Drosophilidae

Among the several parasites and inquilines that are attracted by the rich resources and clean and protective shelter of the Western honey bee *Apis mellifera* nest, none has undergone as profound morphological changes as the apterous and quasi-blind bee louse fly *Braula coeca* (Figure 1A-C). The female lays eggs in honey (not brood) cells, and the hatched larvae eat pollen and wax, where they burrow tunnels in which they pupate without forming true puparia.^4,5^ Following emergence, the adults attach to the body of worker bees, migrating from one individual to another until reaching the queen (Figure 1A). There, they move to the queen’s head, stimulate regurgitation, and imbibe from her mouth honey and nectar.^4,5^ The bee louse fly is considered an inquiline kleptoparasite with potential negative effects on honey bee colony health due to the galleries it makes in bee combs and the facilitation of transmitting serious pathogenic viruses to the bees.^8^

**Figure 1.**
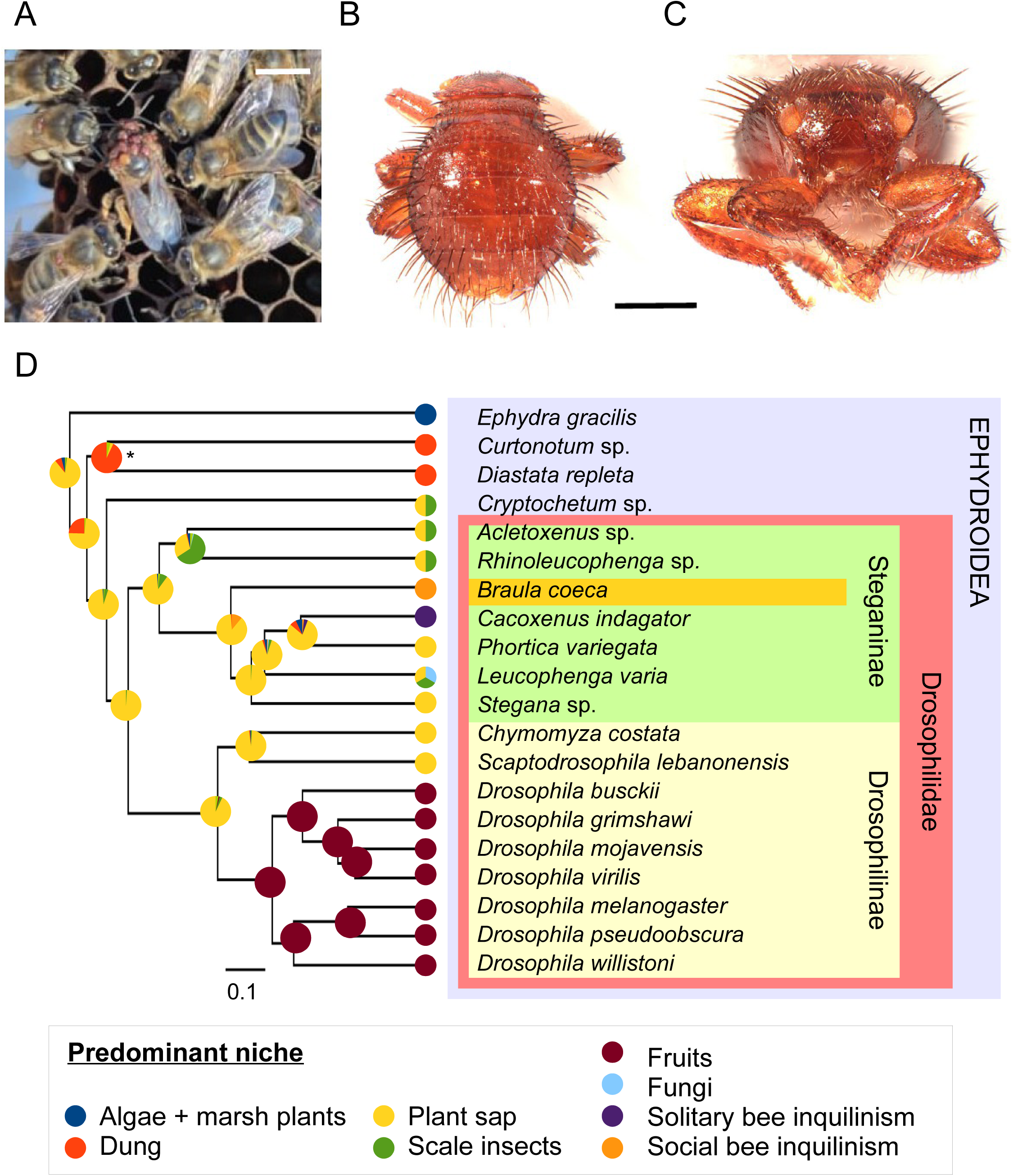
The bee louse fly (*Braula coeca*) is an inquiline of the Western honey bee (*Apis mellifera*) and has likely evolved from a sap-breeding drosophilid associated with scale insects. (A) Tens of *B. coeca* adults preferentially attached to the honey bee queen (© Etienne Minaud). Scale bar = 5 mm. (B) Dorsal view of an adult showing the loss of the wings, halters, and scutum, mesonotum reduction and the legs’ robustness. Scale bar = 0.5 mm. (C) Frontal view of an adult showing the reduction of the eyes and the loss of the ocelli. Scale bar = 0.5 mm. (D) Maximum likelihood phylogeny inferred from 3,100 conserved single-copy proteins (2,557,349 amino acids) showing the position of *B. coeca* (orange) in the subfamily Steganinae (light green) of the Drosophilidae (red). Outgroup species belong to the superfamily Ephydroidea (light blue). All internal nodes had an ultra-fast bootstrap value of 100% except * = 73%. Pie charts at internal nodes indicate the likelihood of ancestral breeding niches inferred from the predominant niches of terminal taxa.

Ever since Réaumur’s first description of the bee louse fly in 1740, and Nitzsch’s creation of the genus *Braula* in 1818 (Document S1),^9,10^ the positioning within the Diptera of the bee louse fly and affiliated species that were classified under the family Braulidae has been puzzling because of its aberrant morphology and unique adaptations to a social host. This family contains seven species belonging to the genera *Braula* and *Megabraula* that are all inquilines to honey bee species of the genus *Apis*. Recent phylogenetic analyses based on a transcriptome assembled from one adult fly and using 1,130 loci interestingly showed *Braula coeca*, the most widespread braulid, to constitute a basal lineage within the Drosophilidae that was sister to four genera of the subfamily Steganinae.^6,7^ To reassess this hypothesis using a larger dataset, we sequenced the whole genome from a pooled sample of 15 unsexed *B. coeca* flies, all collected on Ouessant Island in Western France. We used a hybrid approach to assemble a draft genome using long-read Oxford Nanopore Technology (ONT) and short-read Illumina sequencing (see Methods). Benchmarking Universal Single-Copy Orthologs (BUSCO)^11^ gave a score of 95.8% of the Dipteran conserved single-copy orthologs with 1.3% of duplicated genes. Merqury^12^ estimated an assembly consensus quality value (QV) of 41, which exceeds the recommended threshold of QV40 for reference genome.^13^ We assembled two genomes and one transcriptome of three additional steganine genera. We then built a supermatrix of 3,100 BUSCO genes (2,557,349 amino acids) that included 15 drosophilid species (representative members of the four main radiations in the family),^14^ and 5 species belonging to the superfamily Ephydroidea to which both the Drosophilidae and Braulidae belong (Data S1). The maximum-likelihood phylogenetic analysis of this large dataset reconfirmed the close-relatedness of *B. coeca* to the Drosophilidae. It further showed that it is a full member of the subfamily Steganinae (Figure 1D). The taxonomic priority principle should consider the family Drosophilidae, described in 1856,^15^ a junior synonym for the family Braulidae, described in 1853.^16^ However, the asymmetric size and scientific relevance of the two families argue against such a decision. We, therefore, opt for synonymizing the Braulidae with the Drosophilidae, referring hereafter to *Braula* and *Megabraula* as members of the subfamily Steganinae.

### Inquilinism in the bee louse fly likely evolved from sap breeders associated with scale insects

To gain further insight into the history of the association between *Braula* and *Apis*, we mapped the predominant ecological habitats of ephydroid families on the phylogeny. The ancestral habitat of ephydroids was presumed to be rotting leaf molds.^17^ From this, multiple specializations took place, including the exploitation of aquatic molds (and eventually algae) in the Ephydridae,^18^ Mammal dung in Curtonotidae and Diastatidae,^18,19^ and fermenting vegetables and fruits, sap and fungi, with specialization mostly on yeasts in the Drosophilidae^20^ (Figure 1D). Bayesian reconstruction suggests the ancestral habitat of the Drosophilidae to be tree sap breeding (Figure 1D) with fungus– and fruit-breeding subsequently deriving and predominating in the genera *Leucophenga* and *Drosophila*, respectively. Remarkably, the deepest branches in the Steganinae and the Cryptochaetidae (the closest relative to the Drosophilidae) represent lineages whose larvae are predatory of scale insects and mealy bugs, e.g., *Acletoxenus formosus* and *A. indicus* on aleyrodoids, *Rhinoleucophenga brasiliensis* and *R. obesa* as well as *Cryptochaetidum iceryae* and *C. grandicorne* on coccoids.^21^ In those lineages, adults are often seen to feed on the honeydew produced by the bugs, an abundant sugar-rich substrate sucked from plants’ sap, while larvae take shelter and develop in the waxy secretions of these insects. This dependence on sugary substrate (honeydew) and development in a waxy environment could have predisposed *Braula*’s inquilinism in bee nests.

### The bee louse fly inquilinism is relatively recent

To date *Braula* inquilinism, we inferred a fossil-calibrated phylogeny using 79 single-copy orthologs (63,192 amino acids) in 17 Acalyptrate dipteran and 25 Apocrite Hymenopteran species (see Methods; Data S1; Figure 2A). Five non-ephydroid dipteran species with Ref-Seq assemblies were included in this analysis to correct for tree imbalance^22^. The divergence between *B. coeca* from its closest steganine relatives (1 in Figure 2A: 44.9 [37.8-53.8] million years (myr) [95% confidence interval]) overlapped with the origin of the Apidae (50.12 [42.9-64.5] myr) and with the transition from solitary to subsocial (2: 40.25 [32.4-47.8] myr) and primitively social habits (3: 30.98 [26.3-36.0] myr).^23^ It is possible that the origin of the bee louse fly-apid interactions occurred at sap breeding sites, when early subsocial apids started to gather resin and other plant exudates, as well as scale insects honeydew, and stored them in their nests. As eusociality evolved (4: 23.8 [18.9-28.2] myr), the proportion of resin to secreted wax diminished, and some cells were also used to store nectar and honey for the brood.^24^ A shift from the putatively ancestral dependence on honey and wax produced by scale insects to those produced by bees might have evolved by then. The transition to eusociality in the genus *Apis* required an important division of labor that involved the evolution of pheromonal control of the reproductive capacity of worker females by the queen and the evolution of trophallaxis.^24^ Adaptation of *Braula* to the queen pheromone compounds that have anti-ovarian effects on a wide range of insects, including *Drosophila melanogaster*^25^ and the exploitation of trophallaxis^5^ could not have evolved before the advancement of eusociality (5: 17.1 [11.6-23.0] myr). The evolution of blindness and apterism should have constrained the dispersal of the bee lice, relating their speciation history to that of their hosts. Indeed, only seven bee louse fly species are known, of which five *Braula* species are restricted to the Western honey bee *A. mellifera*, and two *Megabraula* species are restricted to the giant honey bee *A. laboriosa* in the Himalayas.^26,27^ The divergence between these *Apis* species, and presumably between *Braula* and *Megabraula*, is estimated at 6: 5.8 [2.8 –12.0] myr ago. Therefore, the evolution of the bee louse fly inquilinism has likely taken place during the Mid-to Late Miocene period between 5.8 and 17.1 myr ago (Figure 2A). We cannot rule out even a more recent origin if the ancestor of *Braula* or *Megabraula* has shifted from one *Apis* host to another, *i.e.* <5.8 myr ago.

**Figure 2.**
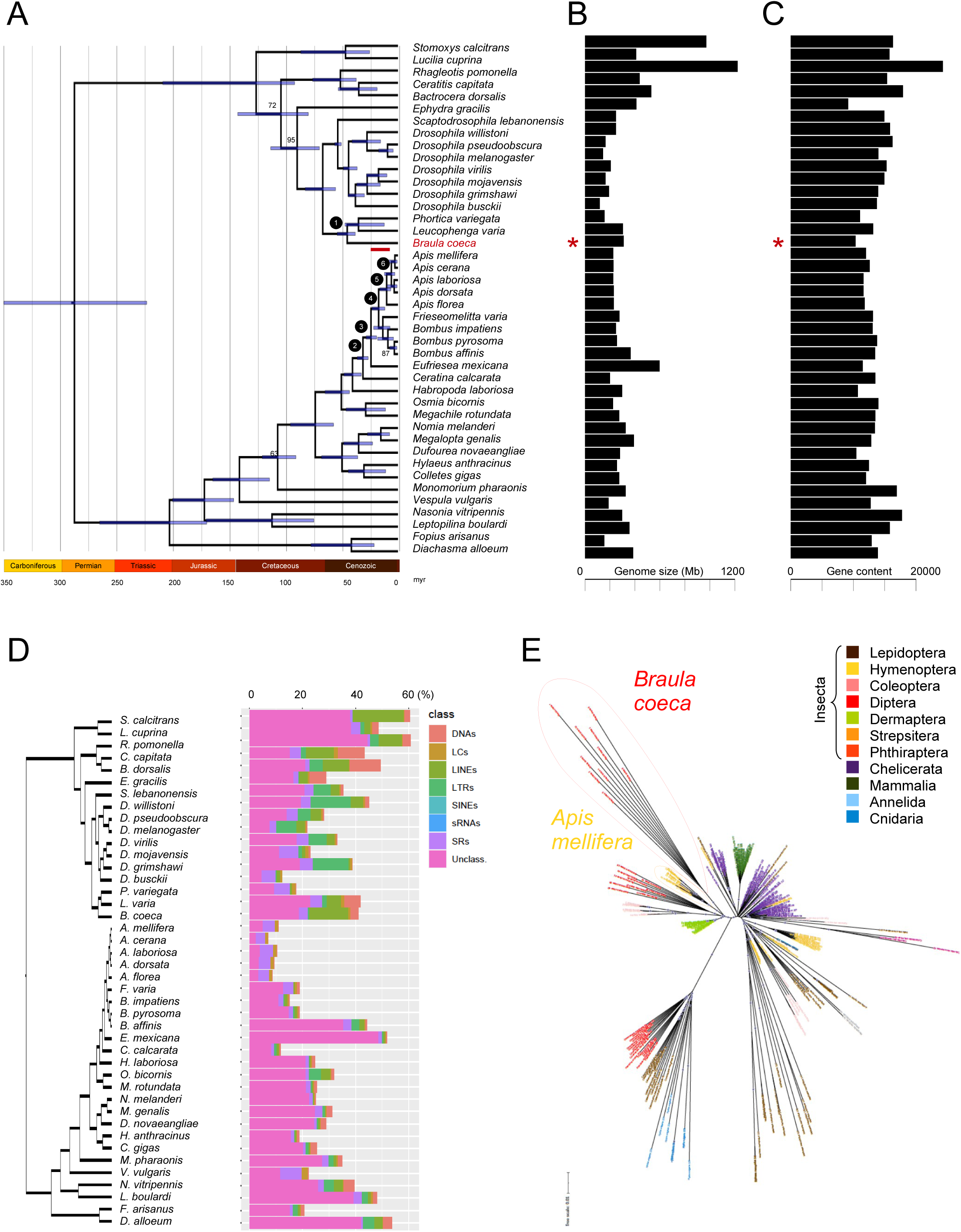
Evolution of the bee louse fly inquilinism, its genome size, gene content, and transposable elements in the bee louse fly with evidence for horizontal transfer between the inquiline and its host. (A) Fossil-calibrated maximum-likelihood phylogeny inferred from 79 conserved single-copy proteins (63,192 amino acids) demonstrating major stages in the evolution of the inquiline and its social host. All internal nodes had an ultra-fast bootstrap value of 100 (except when given), with a blue interval indicating a 95% confidence level of divergence time estimate inferred by MCMCTree. The red bar indicates the likely interval of the origin of the bee louse fly-*Apis* association. Labels 1-6 refer to the major stages mentioned in the text. (B) Genome-size evolution. Red asterisk indicates the estimate for *B. coeca*. (C) Gene content evolution. Red asterisk indicates the estimate for *B. coeca*. (D) Proportions of transposable elements in the genomes of 42 dipteran and hymenopteran species. DNAs = DNA transposons, LCs = low complexity elements, LINEs = long interspersed nuclear elements, LTRs = long terminal repeats, SINEs = short interspersed nuclear elements, sRNAs = small RNAs, SRs = single repeats, and Unclass. = unclassified. (E) Maximum-likelihood phylogeny of *Famar1*-like copies from 38 animal species. Filled circles indicate ultrafast bootstrap values higher than 90%.

### The bee louse fly inquilinism was accompanied by a reduction of gene content but not genome size

Loss of significant portions of genomic and gene contents is a characteristic of obligate parasites specializing on specific hosts or inhabiting extreme environments. For example, the human body louse, *Pediculus humanus*, has one of the smallest genomes and the lowest numbers of genes in insects (108 megabases [Mb] and 10,773 protein-coding genes).^28^ For *B. coeca*, we obtained a final assembly size of 309.35 Mb shared by 2,477 contigs with an N50 of 347,211 bp. This N50 estimate is typical of hybrid genome assemblies obtained using a pooled sample of wild-caught drosophilid flies from species with large genome sizes (> 300 Mb).^29^ No evidence for polyploidy or other endosymbiont that could have biased the genome size estimate was detected (Figure S1). Genome size prediction using k-mers distribution spectra predicted a genome of 308 Mb, concordant with the assembly size (Figure S1). Such a genome size is significantly larger than the remaining drosophilid species (Student’s *t* one-sample test *P* < 1.3 x 10^-4^). Phylogenetic analysis of genome size evolution indicates that the *B. coeca* genome likely retained the size of the ancestral Steganinae, *i.e.* a stronger reduction occurred in the Drosophilinae lineage containing *D. melanogaster* (Figure 2B; Figure S2; Data S1).

To determine the number of protein-coding genes, we used four rounds of Maker^30^ supported by the training of the gene finding and prediction tools SNAP^31^ and Augustus^32^. The annotation, made on the repeat-masked genome, yielded 10,349 protein-coding genes with an Annotation Edit Distance (AED) ≤ 0.5 for 96.4% of our gene models and a Pfam domain found in 83.66% of the proteins. Using the same strategy, we annotated two steganine genomes, namely *Phortica variegata* and *Leucophenga varia*. The annotation yielded 11,067 (BUSCO score = 91%) and 13,160 (BUSCO score = 90.8%) protein-coding genes, respectively. The annotation of the ephydrid *Ephydra gracilis* genome yielded 9,154 protein-coding genes (BUSCO score = 68.9%) (Figure S2). *Ephydra* is particular among Ephydroidea in adapting to hypersaline waters and associated algal flora.^33^ Given the current low knowledge of ephydrid genetics, whether their low gene content is due to their high specialization or an artifact of incomplete annotation, is hard to know. Regardless, the bee louse fly has the lowest number of protein-coding-genes compared to other drosophilids (Student’s *t* one-sample test *P* < 1.5 x 10^-5^) despite having a total genome size that is among the largest genomes in the family. Remarkably, a low gene content is characteristic of bee genomes, compared to ants and wasps, with a remarkable trend of gene reduction within the family Apidae during the evolution of the genus *Apis* (Figure 2C; Figure S2; Data S1).

### Transposable elements (TEs) expanded in the bee louse fly with one element horizontally transferred with the host

The bee louse fly’s large genome size and low gene content suggest an increase in repetitive sequences. RepeatModeler and RepeatMasker analyses^34,35^ indicated that nearly 41.34% of the *B. coeca* genome consists of such sequences, compared to 22.05% and 10.98% in *D. melanogaster* and *A. mellifera*, respectively (Figure 2D). Remarkably, half of the bee louse fly repetitive sequences consisted of long interspersed nuclear elements (LINEs) retrotransposons (14.94%). While LINEs are usually among the most abundant transposable elements after LTRs within the Drosophilidae,^36^ their values did not exceed what was found in *B. coeca* (we found the highest percentage in *Leucophenga varia* with 5.54%). It is at present unclear what factors influence the diversity of transposable elements (TEs) landscapes among eukaryote species^37^. Nonetheless, this difference means that whereas the bee louse fly has likely retained the ancestral large genome size of the Drosophilidae, its TEs constitution has largely evolved.

Because host-parasite relationships have repeatedly been invoked as a factor that may favor horizontal transfer of TEs,^38^ we searched for evidence of such transfers between *B. coeca* and *A. mellifera* (Document S2). We found one TE, a DNA transposon *Famar1-like* element previously described in the earwig *Forficula auricularia*^39^ that belongs to the Tc1-mariner superfamily. This element showed a high similarity between *B. coeca* and *A. mellifera* but was absent in all other drosophilid species for which a genome is available in GenBank, which is highly suggestive of an acquisition through horizontal transfer (Figure S3). Indeed, phylogenetic analysis of multiple copies of this TE extracted from 37 widely divergent animal species (Figure 2E) supported a direct transfer event between *B. coeca* and *A. mellifera*, although the directionality of the transfer cannot be inferred since the elements from the two species form mutually-exclusive monophyletic clades. Remarkably, all elements found in the genomes of four *A. mellifera* subspecies, including *A. m. carnica*, *A. m. caucasia*, *A. m. mellifera* and *A. m. ligustica*, formed an exclusively monophyletic clade. The transfer time between *B. coeca* and *A. mellifera* has likely preceded the dispersion of this element among the subspecies or even their differentiation 0.77 myr ago^40^ if the element was ancestral in *A. mellifera*. On the other hand, we did not find any trace of this element or any other related element in any other *Apis* species, indicating that the maximal time of horizontal transfer likely does not surpass 2.73 [0.70-8.9] myr ago, *i.e.* the time of divergence between *A. mellifera* and its closest-relative *A. cerana* (Figure 2A). The tight ecological connection between the bee louse fly and its host may have favored this transfer, as was suggested for blood-or sap-sucking insects.^41,42^

### Gene families with excess losses show striking cross-order parallelism

Despite their deep divergence, we tested whether parallel changes could explain the reduction of protein-coding genes in both the honey bee and the bee louse fly. We used OrthoFinder^43^ to cluster orthologous proteins from the 25 hymenopteran and 17 dipteran species. We identified 19,010 orthogroups. Of these, 935 showed significant size evolution among the 42 species when analyzed using CAFE5^44^ and after applying an error model that accounted for misassemblies and misannotations. To classify those orthogroups into functional categories, we extracted groups that contained *D. melanogaster* orthologs for which a molecular function, *i.e.* a gene group, was assigned in the Flybase database,^45^ e.g., the 60 odorant receptors of *D. melanogaster* clustered in 16 orthogroups. Of 1,078 gene groups, 163 significantly deviated from the birth-death model estimated by CAFE5.

Thirty-nine gene groups had significant losses in the bee louse fly. Most of these groups showed a striking parallelism with *Apis* in particular and hymenopterans in general (Figure 3). The most significant groups were those involved in the chemical detection of taste (gustatory receptors and divergent ionotropic receptors) and odors (odorant receptors and odorant binding proteins). The remaining groups included those involved in metabolism and/or immunity, such as C-type lectins (recognition), CLIP serine proteases (signaling), serpin (signaling), Dorsal (signaling), lysozyme (other), as well as in detoxification, such as cytochrome P450, GST-C, carboxylases, ABCC transporters, acyl-co, and alcohol dehydrogenase. Indeed, bees have evolved a reduced repertoire of immunity and detoxification genes, likely due to the evolution of social behavior and their life in an overprotective and clean shelter, *i.e.* the nest.^46,47^ Cytochrome P450 genes are more expressed in foraging workers than in the castes that remain in the nest (*i.e.* the queen and nurse workers).^48^ This is also the case of alcohol dehydrogenase,^49^ an emblematic enzyme in *D. melanogaster* that was lost in *B. coeca*. The reduction of peptidases in both the honey bee and the bee louse fly could also be due to the low protein content of some of their food, *i.e.* nectar and honey. We also noted an underrepresentation of chitin-binding domain proteins and chitinases in the bee louse fly and the honey bee. Cuticles could act as barriers against environmental toxins, which may not be highly encountered in the nest. Remarkably, *B. coeca* is unique among cyclorrhaphan dipterans as its pupa, similarly to the honey bee’s,^50^ is contained in the unmodified cuticle of the third instar larva, and no sclerotized puparium is formed.^4,5^

**Figure 3.**
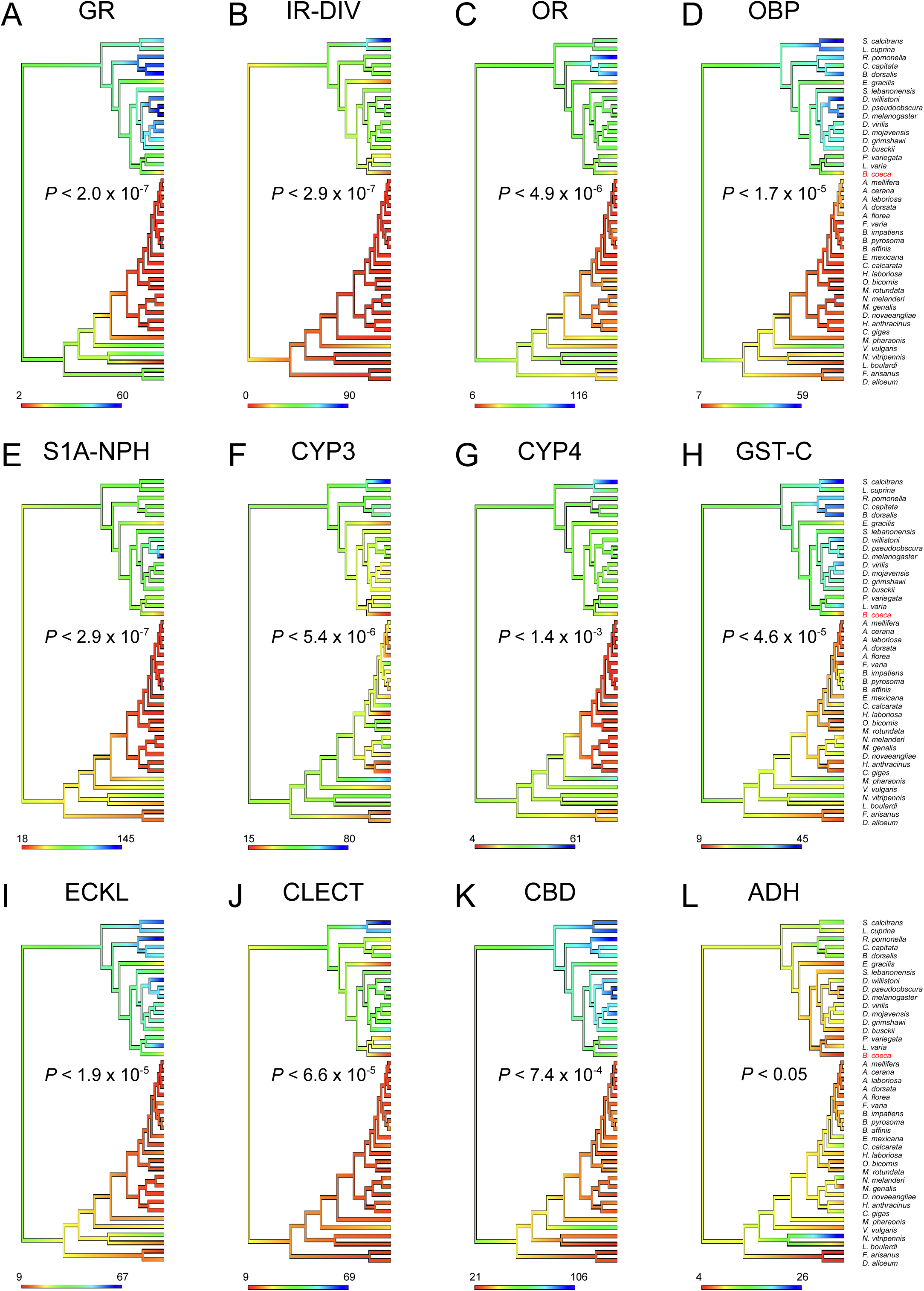
Deep convergence in ecologically-relevant gene families evolution between the bee louse fly and its hymenopteran host. A sample of significantly evolving gene families in *B. coeca* lineage as inferred by CAFE5 from functionally-grouped orthogroups defined by OrthoFinder (see text). For each panel, the species tree from Figure 2 is given with branch colors reflecting inferred ancestral family size reconstruction using fastanc in phylotools. CAFE5 *P*-values are noted for each panel. (A) GR = gustatory receptors. (B) IR-DIV = divergent ionotropic receptors. (C) OR = odorant receptors. (D) OBP = odorant-binding-proteins. (E) S1A-NPH = S1A non-peptidase homologs. (F) CYP3 = Cytochrome P450 CYP3 clan. (G) CYP4 = Cytochrome P450 CYP4 clan. (H) GST-C = Cytosolic glutathione S-transferases. (I) ECKL = Ecdysteroid kinase-like. (J) CLECT = C-type lectin-like. (K) CBD = Chitin binding domain-containing proteins. (L) ADH = Alcohol dehydrogenases.

### Honey and wax feeding drove the loss of almost all bitter-tasting gustatory receptors

The two most significantly evolving gene families in the bee louse fly, *i.e.* gustatory receptors (GRs) and divergent ionotropic receptors (IR-DIVs), allow the detection of soluble cues (Figure 3A-B). There are 60 GRs in *D. melanogaster*, of which 9 and 49 receptors respond primarily to sweet and bitter tastes, respectively, and 2 receptors respond to carbon dioxide (CO_2_).^51^ The three categories clustered into 35 orthogroups (Figure 4A), whose phylogenetic analysis indicates that the ancestral drosophilid repertoire consisted of 6 sweet, 29 bitter, and 2 CO_2_ GRs assuming functional conservation of gustatory categories (Figure 4A). We identified 11 GRs in the bee louse fly with no duplications using InsectOR^52^ and manual curation. These GRs could be classified according to their *D. melanogaster* orthologs into 2 sweet, 7 bitter, and 2 CO_2_. That means that the *D. melanogaster* lineage disproportionally evolved more bitter receptors from the ancestral repertoire, whereas *B. coeca* disproportionally lost bitter receptors (Figure 4A). InsectOR inferred the number of GRs in the steganine species *L. varia* and *P. variegata* to be 21 and 26, respectively, further confirming that *B. coeca* has lost a significant portion of the ancestral GR repertoire (Figure S4). Honey bees have only 11 GRs, of which 7 are orthologous to sweet *Drosophila* GRs.^53^ This is likely due to the bees’ strong diet reliance on sweet floral nectars and honey.^54^ The loss of *B. coeca* bitter GRs and its retention of 2 ancestral sweet receptors is a strong convergence with its host.

**Figure 4.**
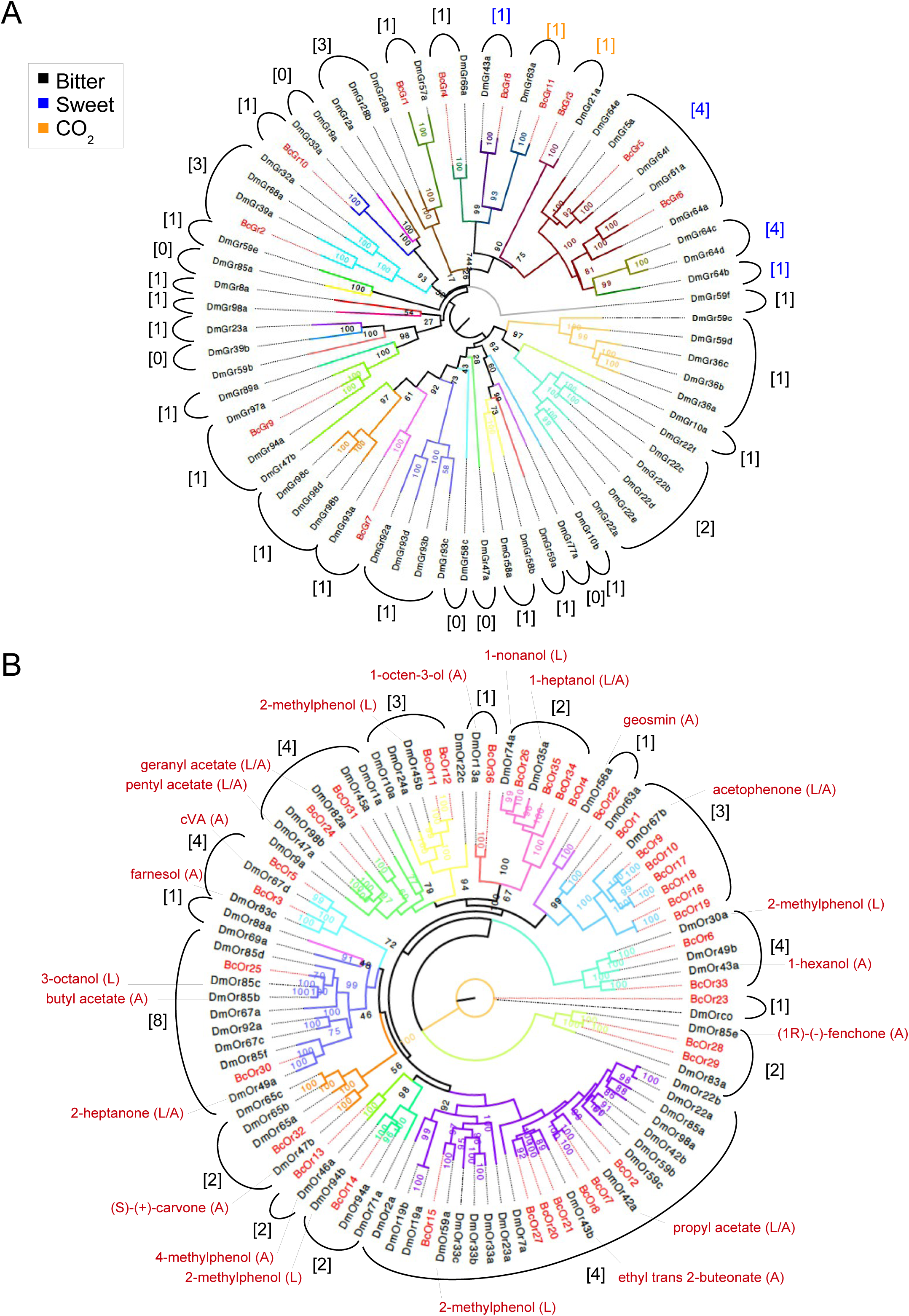
Evolution of chemosensory receptors gene families in *Braula coeca* and *Drosophila melanogaster*. (A) Maximum-likelihood phylogeny of gustatory receptors (GRs) with main taste categories color code given in a frame. (B) Maximum-likelihood phylogeny of odorant receptors (ORs) with main ligands for each *D. melanogaster* receptor given in dark red. L = larva and A = adult expression. For A and B, ultra-fast bootstrap values are given above nodes. Branches are colored according to orthogroups defined by OrthoFinder for 42 dipteran and hymenopteran species. Numbers in broken brackets before each orthogroup reflect the presumed ancestral gene content inferred by phylotools.

Ionotropic receptors are another major class of chemoreceptors. They are divided into antennal IRs, which are conserved across insects and are most likely involved in olfaction, and divergent IRs (IR-DIVs), which evolve rapidly and are mostly involved in the taste perception of carboxylic and amino acids. Only divergent IRs showed a significant loss in *B. coeca* (Figure 3B). However, our knowledge about the function of the 42 *D. melanogaster* IR-DIVs is still limited.^55^ We inferred the ancestral IR-DIV drosophilid repertoire to contain 29 receptors, of which only 9 were retained in *B. coeca*. Remarkably, whereas we found almost no direct orthologs between Diptera and Hymenoptera for IR-DIVs (Figure 3B), bees are known to have few IRs in general^56^ pointing to another possible taste convergence between the bee louse fly and its host.

### One-fifth of ancestral odorant receptors was lost, including one receptor that is involved in anti-ovarian response in Drosophila melanogaster

Odorant receptors (ORs) are essential to detect volatile chemical cues from the environment. This family has expanded in the honey bee to reach 170.^57^ However, only 9 of the honey bee genes have orthologs with *D. melanogaster*, and phylogenetic analysis indicates that this common OR repertoire has been gradually reduced during the evolution of *Apis* (Figure 3C). The 60 ORs of *D. melanogaster* clustered within 16 orthogroups (Figure 4B). We inferred the ancestral drosophilid OR repertoire to contain 44 ORs, with at least one representative for each orthogroup (Figure 4B). We identified in *B. coeca*, following InsectOR^52^ and manual curation, 35 ORs in addition to *Orco*, *i.e.* one-fifth of the ancestral repertoire was lost. The number of ORs was 50 and 51 in the two closely related steganine species *L. varia* and *P. varia*, respectively. *Braula* ORs were direct orthologs to 18 genes in *D. melanogaster* (Figure 4B). Judging from the response of these orthologs to different volatiles in *D. melanogaster* as curated in the DoOR database^58^ and assuming potential conservation of function, the retained bee louse fly ORs may respond to compounds produced by honey bee workers in a defense context (e.g., 1-hexanol, farnesol, 2-heptanone),^59^ and/or of floral, pollen and nectar aromas, such as acetophenone and benzaldehyde, a major volatile of honey.^60,61^ Two cases of tetraplications were observed. One case involved three recent duplications of genes orthologous to DmOr67b, a gene that is highly responsive in *D. melanogaster* to both acetophenone and 1-hexanol. The second case involved three successive duplications of a gene orthologous to DmOr74a, which responds in *D. melanogaster* larvae to 1-nonanol and 1-heptanol, the latter being a major brood volatile,^10^/23/23 11:02:00 AM and 1-hexanol, a component of the alarm pheromone.^62^ Of these three duplications, two were unique to *B. coeca* compared to its closely-related steganine species (Figure S5). Low concentrations of isopentyl acetate, the main component of the alarm pheromone, released by unstressed workers at the nest entrances attract the parasitic nest beetle *Aethina tumida*,^63^ suggesting that the detection of the host odors could be a common strategy among phylogenetically distant inquilines and parasites of social insects.

Whereas major molecular convergences could exist between the inquiline and its social host, divergent strategies to adapt to the eusocial lifestyle requirements are still needed. In honey bees, colony cohesion is driven by the volatile queen’s mandibular pheromone (QMP), which “sterilizes” the bee workers.^64^ This pheromone elicits an anti-ovarian response in other insects, including *D. melanogaster*.^25^ An RNA interference (RNAi)-screen identified DmOr49b, DmOr56a, and DmOr98a to be potentially involved in the detection of the QMP compounds and the suppression of fecundity.^25,65^ A *sine qua non* condition for a drosophilid to reproduce in a bee nest would, therefore, be to lose those receptors or to modify their response or effect. We found that the bee louse fly does not have an ortholog for DmOr98a, a receptor specific to the genus *Drosophila* (Figure S5). The bee louse fly has a pseudogene, orthologous to DmOr49b, that was InsectOR identified. Orthologs of this *D. melanogaster* receptor are present and complete in all dipteran species, including *L. varia* and *P. variegata* (Figure S5). The bee louse fly had a receptor that we called BcOr22, which was orthologous to DmOr56a (Figure 4B). This last receptor is narrowly tuned in many *Drosophila* species to a single component, the mold volatile geosmin, whose perception also inhibits oviposition in *D. melanogaster*,^66^ pointing to a possible conserved role in reproduction. Therefore, further functional analyses of the response of candidate ORs to various QMP compounds are required in both *D. melanogaster* and *B. coeca* to understand how modifications of these genes in *B. coeca* might have facilitated the evolution of the bee louse fly inquilinism.

### Blindness and life in a dark nest were accompanied by the loss of multiple rhodopsins

The species Latin name of the bee louse fly refers to the assumption that it was blind due to the reduction of the eye size and the loss of the ocelli. In agreement with reduced vision in the bee louse fly, we found only two out of the seven rhodopsin genes, which are responsible for colored vision and positive phototaxis in *D. melanogaster* and which were all present in the ancestral drosophilid repertoire. *D. melanogaster* orthologs of the *Rh1* and *Rh6* genes are expressed in the ommatidia and are sensitive to light.^67^ The role of these opsins in light detection, despite the absence of ommatidia in the bee louse fly is unclear. Remarkably, *Rh1* and *Rh6* are structurally required in mechanosensory bristles to control larval locomotion.^68^ They also detect temperature.^69^ Therefore, the retention of these rhodopsins in the bee louse fly could mainly be due to their unconventional functions. On the other hand, the rhodopsin *Rh2*, which is exclusively expressed in the ocelli and used for horizon detection in *D. melanogaster* ^70^, is among those lost in the bee louse fly, in agreement with the loss of the ocelli. Regression of the visual system and its underlying opsin genes is common in animals inhabiting dark environments, such as fossorial mammals^71^ and cavefishes,^72^ representing a major example of deep convergences.

### Apterism was not accompanied by the loss of major wing development genes

Small size, loss of wings, and the evolution of strongly clinging legs are all morphological changes that could prevent the honey bees from getting rid of the bee lice.^73^ All these potential adaptations are convergent with ectoparasitic true lice, and for some, such as apterism, represent major recurrent changes that have responded to distinct pressures throughout the history of insects.^74^ We found intact most of the main wing development genes whose mutations severely reduce the wing in *D. melanogaster*, such as *wingless*, *apterous*, or *vestigial*. This means that the major morphological changes more likely resulted from regulatory changes of these core genes or modifications of other genes. Future developmental studies, specifically comparing the expression of wing and leg morphogenic genes between the bee louse fly and *D. melanogaster*, will definitively help shed light on the transcriptomic shifts underlying the major morphological changes of the bee louse fly.

## Conclusion

That the enigmatic bee louse fly is indeed a drosophilid, a lineage within the most investigated insect family with more than 150 fully sequenced genomes, is undoubtedly one of the most exciting discoveries in dipteran phylogeny. How could a fly with an ancestral drosophilid genome become ecologically adapted to bees and morphologically similar to lice? Our results show that a mosaic of deep convergences at the genomic level underlies the relatively recent and dramatic changes of the bee louse fly to nest inquilinism. This mosaicism involved deep convergences with the host, mostly in genes likely involved in immunity, detoxification, and chemical perception, as well as convergences with general features of fossorial animals in the visual systems. Future developmental studies may elucidate whether general morphologies, such as apterism and leg modifications, could also be shared between *Braula* and other ectoparasites. Due to its genetic relatedness to *Drosophila* and ecological association to *Apis*, two major laboratory models, the new genomic resources presented here can help establish the bee louse fly as a promising model to address questions related to deep convergences that are difficult to approach in multiple highly specializing animals.

## Acknowledgments

We are very grateful to Marcus Stensmyr for insightful comments on an early draft of the manuscript, the *Association Conservatoire de l’Abeille Noire Bretonne* (A.C.A.N.B.) for help collecting *B. coeca* flies, Etienne Minaud for photographic images of *B. coeca* and Héloïse Muller for help with the genome annotation pipeline, Christian Cheminade and Michael Lang for help translating early literature on *Braula*. *Braula* genome sequencing was funded by a grant from Université Paris-Saclay (ADAPAR) to HB.

## Author contributions

Conceptualization: H.B. and A.Y.; Investigation: H.B., H.L., N.D., D.O., C.P., J.F., C.G., and A.Y.; Resources: H.L. and L.G.; Writing – Original draft: H.B., H.L., D.O., J.F., C.G., and A.Y.; Writing – Reviewing & Editing: H.B., J.C., C.G., F.M.P., F.R., J.C.S., and A.Y.; Visualization: H.B., J.F., C.G., and A.Y.; Supervision: H.B.; Funding Acquisition: H.B.

## Declaration of interests

The authors declare no conflicts of interest.

## Declaration of generative AI and AI-assisted technologies in the writing process

During the preparation of this work the authors used Grammarly (Grammarly Inc.) in order to improve language and readability. After using this tool/service, the authors reviewed and edited the content as needed and take full responsibility for the content of the publication.

## Methods

### Lead contact

Further information and requests for resources and reagents should be directed to and will be fulfilled by the lead contact, Héloïse Bastide (<u>heloise.bastide@universite-paris-</u> <u>saclay.fr</u>). Raw sequence data are deposited on NCBI Sequence Read Archive (SRA) under Bioproject PRJNA1000103. Perl scripts are available on Github depository at https://github.com/AmirYassinLab/Supermatrix and https://github.com/AmirYassinLab/OG2GG. Commands for all programs are available on Github depository at https://github.com/hbastide/Braula_genome_analysis. Genome assemblies and all data associated to this study are deposited in Figshare and will be made available upon publication.

### Sample collection and genomic library preparation

Samples of *Braula coeca* were collected from honey bee colonies on the Island of Ouessant in France and kindly provided to us by the *Association Conservatoire de l’Abeille Noire Bretonne* (A.C.A.N.B.). Genomic DNA was extracted from 15 unsexed individuals conserved in alcohol using the Nucleobond AXG20 kit and buffer set IV from Macherey-Nagel (ref. 740544 and 740604, https://www.mn-net.com, Düren, Germany).

### Genome sequencing and assembly

We used a hybrid approach to assemble a draft genome of *B. coeca* using both long-read Oxford Nanopore Technology (ONT) and short-read Illumina sequencing.^75^ Before nanopore sequencing, a size selection was conducted on the DNA using the SRE XS from Circulomics (https://www.circulomics.com/, Baltimore, Maryland, USA). The SQK-LSK110 kit from ONT (https://nanoporetech.com/)^76^ was then used to prepare the samples for nanopore sequencing following the manufacturer’s protocol. The library was loaded and sequenced on an R9.4.1 flow cell (ref. FLO-Min106) for sequencing. Raw data were basecalled using Guppy v5.0.11 and the “sup” algorithm. The ONT raw data size was 4.4 Gb in 1,399,323 reads (mean read length 3,146 kb, longest read of 123.3 Mb), with an N50 of 4,677 kb. Phred scores ranged from 8 to 18, with a median of 13, as assessed by PycoQC.^77^ Illumina paired-end sequencing was performed by Novogene Company Limited (https://en.novogene.com, Cambridge, UK) on the same DNA sample. The Illumina sequencing produced 119,719,537 paired 150 bp reads. Phred scores averaged 36 per read as analyzed by FastQC (http://www.bioinformatics.babraham.ac.uk/projects/fastqc/). We used MaSuRCA v4.0.3^78^ to produce the hybrid assembly of our genome using the Cabog assembler. We obtained a final assembly size of 309,35Mb in 2477 contigs, with a N50 of 347,227 bp. The completeness of the assembly was estimated to 95.8% with BUSCO v5.0.0 on the *diptera_odb10* dataset (C:95.8%[S:94.5%,D:1.3%],F:0.7%,M:3.5%,n:3285), and to 93.6% using Merqury.

### Estimation of genome size and endosymbionts detection

K-mers frequencies within short-read data were obtained with KMC 3.^79^ Genome size and ploidy were inferred using GenomeScope v2.0 with k-mer size = 21 and Smudgeplot.^80^ Contig taxonomy was performed using Blobtools^81^ with Diamond as search engine^82^ against the UniProt database using a local copy of the NCBI TaxID file for the taxonomic assignation of the best hit. Minimap2^83^ was used for read mapping.

### Genome annotation

The *B. coeca* genome was annotated using Maker v2.31.10,^30^ following Muller et al.’s^84^ protocol, wherein multiple rounds of Maker supported by the training of the SNAP v.2006-07-28^31^ and Augustus v.3.3.3^32^ gene finding and prediction tools, were conducted. RepeatModeler v2.0.1 was first used to identify the repeat-enriched regions that were masked by RepeatMasker v4.0.9 as implemented in Maker. Proteomes of five *Drosophila* species, namely *D. innubila*, *D. albomicans*, *D. bipectinata*, *D. melanogaster*, and *D. virilis* were obtained from NCBI and used to guide the annotation. Protein-Protein BLAST 2.9.0+^85^ (-evalue 1e-6 –max_hsps 1 –max_target_seqs 1) was then used to assess putative protein functions in *B. coeca* by comparing the protein sequences given by Maker to the protein sequences from the annotated genome of *D. melanogaster*. The completeness of genome annotation was assessed using BUSCO at each round and the round with the highest score was retained.

### Phylogenomic analysis of the Ephydroidea

Besides our *B. coeca* assembly, we obtained from NCBI repository genome assemblies for 12 species, transcriptome shotgun assemblies (TSA) for four species, and sequence read runs (SRR) for three species (Data S1). Paired-end DNA raw data of two species, namely *Rhinoleucophenga* cf. *bivisualis* and *Cacoxenus indagator* were assembled using MaSuRCA with default parameters. The transcriptome of *Acletoxenus* sp. was assembled using Trinity software package^86^ on the Galaxy Europe website^87^ following standard protocol^88^. BUSCO v.5.0 was used to assess the completeness of those assemblies and to extract single-copy BUSCO genes for all species. Protein sequences of 3,100 single and complete BUSCO genes were aligned using MAFFT^89^ and concatenated into a single supermatrix (2,557,349 amino acids). A maximum-likelihood (ML) phylogeny was then inferred for the supermatrix using IqTREE 2^90^ with 1,000 ultrafast bootstrap iterations^91^ and using the JTT+R substitution model inferred by ModelFinder^92^ implemented by IqTree.

### Reconstruction of the ancestral ecological niches

For each of the 20 ephydroid species we obtained a predominant ecological niche from the taxonomic literature.^17,20,21,93^ Eight predominant niches were coded as discrete traits, and the Multistate program of the BayesTraits v.4 package^94^ was used under the Reverse Jump MCMC model with 1,010,000 chain iterations and a burnin sample of 10,000.

### Phylogenomic analysis of Diptera and Hymenoptera

The second phylogenomic analysis involved, besides *B. coeca*, 25 Hymenopteran and 16 Dipteran species for which an assembly can be downloaded from the NCBI Genome repository (Data S1). Protein sequences for all species but three, namely *Leucophenga varia*, *Phortica variegata*, and *Ephydra gracilis*, were obtained from NCBI. For these three species, we used the same four-round annotation procedure that we used for *B. coeca* to identify protein-coding genes and translate their sequences. We used BUSCO to assess the completeness of all annotated and downloaded genomes and their corresponding assemblies. OrthoFinder^43^ was used to generate protein sequences of protein-coding-genes of the 42 species and to cluster these sequences into orthogroups. Only the longest isoform (*i.e.* the primary transcript) was used for genes with multiple isoforms. 79 orthogroups contained a single copy ortholog from each species, and their protein sequences were aligned using MAFFT and concatenated into a single supermatrix (63,192 amino acids). A maximum-likelihood (ML) phylogeny was then inferred for the supermatrix using IqTREE 2^90^ with 1,000 ultrafast bootstrap iterations^91^ and the JTT+R substitution model inferred by ModelFinder^92^ implemented by IqTree.

MCMCTree^95^ was used to date the inferred ML trees based on recently published fossil-calibrated phylogenies. First, two time points were obtained for the 42-species phylogeny. These included the divergence between ants and bees between 90-120 myr ago^96^ and between *Scaptodrosophila* and *Drosophila* between 50-56 myr ago,^97^ with a maximum root age for the ancestor of Hymenoptera and Diptera at 344 myr ago.^96^

### Genome size and gene content evolution

Genome size and gene content (number of OrthoFinder generated protein-coding-genes after retaining the longest isoform for genes with multiple transcripts) inferred for each of the 42 Dipteran and Hymenopteran genomes were mapped on the phylogenetic tree, and values at the ancestral nodes were inferred and visualized using the fastAnc command in the R package Phylotools v0.2.2.^98^

### Transposable elements annotation and transfer

Transposable elements were identified in *B. coeca*, *D. melanogaster*, and *A. mellifera* following a two-step protocol. First, we used RepeatModeler v2.0.1^34^ with default parameters to generate a *de novo* library of repetitive regions. RepeatMasker v 4.0.9^34^ was then run with the newly generated library and the options –a (create a .align output file) and –s (slow search; more sensitive) to create a summary of the families of transposable elements found in each reference genome along with the percentage of the genome they represent. Horizontal transfer analysis protocols of the *Famar1*-like element are given in Document S2.

### Gene family evolution

We used CAFE v. 5^44^ to model and infer gene family evolution. We conducted CAFÉ5 using an error model on the 19,011 orthgroups generated by OrthoFinder and using the time-calibrated phylogenetic tree of the 42 Dipteran and Hymenopteran species. We also grouped OrthoFinder orthogroups into functional gene groups using the customized perl script OG2GG.pl (https://github.com/AmirYassinLab/OG2GG). The script extracts and groups orthogroups containing *D. melanogaster* orthologs that were assigned to large functional gene groups in Flybase^45^ (“gene_group_data_fb_2023_02.tsv”). CAFE5 was then run on the gene groups’ gene counts using a single birth rate and the error model to correct for possible assembly and annotation errors.

### Chemosensory superfamilies evolution

To curate *B. coeca* gustatory receptors (GR) and odorant receptors (OR) genes, we queried *D. melanogaster* GRs and ORs protein sequences on *B. coeca*, *L. varia* and *P. variegata* assemblies using Exonerate ver. 2.2^99^ with option –maxintron 2000 and –p pam250. The output, along with the assembly, were fed to InsectOR^52^ with option 7tm_7 and 7tm_6 activated for GR and OR analyses, respectively. From the output files, we extracted 300-500 amino acids-long complete sequences with 7tm_7 or 7tm_6 motif detected and with start codon present and no internal stop codon, *i.e.* pseudogenes excluded. Protein sequences were analyzed using Miniprot^100^ to produce an exhaustive gff file with exact genomic coordinates for each species. Protein sequences were aligned using MAFFT and a maximum-likelihood phylogenetic tree for each family using IqTREE 2 with the same options as the phylogenomic analysis. The literature was reviewed to classify GRs into bitter, sweet, and CO_2_ categories^101^ and identify volatile ligands eliciting the strongest response in odorant neurons in *D. melanogaster*.^58^ Because CAFE5 inferred ancestral counts for orthogroups with significant deviation only, we estimated and visualized ancestral counts for each orthogroups of these two families using FastAnc command in the R package Phylotools.

## List of supplementary materials

**Data S1.** List of all genome assemblies and annotations downloaded or generated in this study with data on genome size and gene content.

**Figure S1. Features of *Braula coeca* genome assembly**

(A) Smudgeplot comparing the sum of heterozygous kmer pair coverages (A+B) to their relative coverage of the minor kmer (B/A+B). The hottest smudge corresponds to expected diploid kmer pairs (AB).

(B) Kmer profile spectrum estimating the length of the genome at 307,538,118 bp generated by GenomeScope2.

(C) Square-binned blob plot showing the distribution of assembly scaffolds on GC proportion and coverage axes. Squares within each bin are colored according to taxonomic annotation and scaled according to total span.

(D) ReadCovPlots generated by Blobtools visualising the proportion of reads of *B. coeca* that are unmapped or mapped and showing the percentage of mapped reads by taxonomic group, as barcharts.

**Figure S2. Evolution of genome size and gene content and completeness of assembly and annotation of 42 dipteran and hymenopteran genomes**.

(A) Genome-size evolution and genome assembly completeness inferred by BUSCO across 42 dipteran and hymenopteran species.

(B) Gene content evolution and genome annotation completeness inferred by BUSCO across 42 dipteran and hymenopteran species.

Branch colors in A and B reflect inferred ancestral reconstructions using fastanc in phylotools. CS = complete and single-copy BUSCOs; CD = complete and duplicated BUSCOs; F = fragmented BUSCOs; M = missing BUSCOs.

**Figure S3. Horizontal transposon transfer between *Braula coeca* and *Apis mellifera*.** Comparison of Famar1-like transposon synonymous distance (dS) and orthologous gene synonymous distances between *Braula coeca* and *Apis mellifera*. The red line indicates the 0.5% quantile (=1.76) of the distribution of orthologous gene dS calculated over 2,179 codon alignments. The distribution is bimodal, with genes having highly saturated dS values showing a peak centered on 9.99 and highly conserved genes showing less saturated dS values showing another peak around 2.5. The *Famar1*-like dS (green line, = 0.12) was calculated over the transposase open reading frame of one copy of the element extracted from the *A. mellifera* genome and another copy from the *B. coeca* genome.

**Figure S4.** Maximum-likelihood phylogeny of gustatory receptors (GRs) from *D. melanogaster*, *B. coeca*, *L. varia* and *P. variegata*.

**Figure S5.** Maximum-likelihood phylogeny of odorant receptors (ORs) from *D. melanogaster*, *B. coeca*, *L. varia* and *P. variegata*.

**Document S1** – Translation of early works on the bee louse fly.

**Document S2** – Analysis of horizontal transfer of transposable elements between *B. coeca* and *A. mellifera*.

## Notes

### Competing Interest Statement

The authors have declared no competing interest.

### Summary of Updates

Additional phylogenetic and gene family evolution analyses

