## Supplementary material for "The genome of the blind bee louse fly reveals deep convergences with its social host and illuminates Drosophila origins": Document S1

**Supplementary text 1 – Translations of early descriptions of *Braula coeca* by Réaumur (1740, p. 711-713)**^1^ **and Nitzsch (1818, p. 286-287 and p. 314-316)**^2^

1. Translation from French to English of the first description of the bee louse by René-Antoine Ferchault de Réaumur (1740, p. 711-713)^1^

A small insect attacks the bees themselves, sucking them for food. They have been given to a species of louse not found on other flies. Young bees don't have them; it's only the old ones, and the old ones in certain hives, that are subject to this vermin. Usually only one can be found on each bee; and to see it, you don't have to look very hard. It is reddish, about the size of the head of a very small pin; it almost always stands on the corselet [thorax]; one would be inclined to take it for a small grain of raw wax which would have remained attached to it: but, when examined with even a weak magnifying glass, it cannot be mistaken; one can distinguish most of its parts very well; its body appears shiny and scaly, as are the six legs which support it. If we use a strong magnifying glass, we can see a large number of hairs on its scaly envelope. There is no head shape at the front; the tip seems to be cut off squarely, and this, because it curves underneath; and this curved portion decreases in size, ending in a fine point, which is undoubtedly the tip of the proboscis. At the top, the curved part has a fairly high tubercle on each side; these two tubercles are presumably the insect's eyes. After the anterior part, there are three well-marked rings [segments], from each of which a pair of legs extend. You have to look for the separations from the other rings on the body to see them; but they are most noticeable on the belly side. The foot at the end of each leg forms a kind of pallet bordered by at least three or four hooks. It's a pleasure to see how the hooks on each foot cling to the bee's hairs, which support the little animal without bending under the weight. I have often found it near the fly's neck, near the origin of its wings, and sometimes near that of some leg. I don't believe its proboscis is capable of piercing the scales that cover the bee's corselet; but it can get into joints where flexibility being needed, the scale must be missing.

We don't have a good idea of the hives from which most of the flies have these lice, and perhaps we're right, because it's more common to find them on the flies of old hives than on those of new hives; they've had more time to multiply; but do they really do much harm to the flies? That's what we don't really know, at least it seems certain that they don't cause them much pain, or even that they don't worry them; for, although it may not be as easy for the fly to get one of its legs over its corselet, as over some other part of its body ; and although this is perhaps what determines the louse to place itself there, it is often in places where one of the fly's legs can be carried, and from where it could cause it to fall, and where it is nevertheless allowed to remain still. Nevertheless, these little insects have been regarded as very harmful to bees. There have been taught ways of killing them, which I don't think are quite certain. One of the most vaunted remedies to rid bees of them is to sprinkle them with urine, to throw some on them in the hive with a kind of brush; but urine does not seem to me to be as harmful to these lice as was thought; and very few will get wet with it. Another remedy, because there are remedies to choose from for bee diseases, as there are for ours, is to sprinkle them with brandy, and another is to smoke them.

2. Translation from German to English of the description of the genus *Braula* by Christian Ludwig Nitzsch (1818, p. 286-287)^2^

C. Appendix of the epizootic Diptera;

The Braula, a parasitic insect of the honey bee, very different from Pediculus apis auctt., and however very different from all two-winged [dipteran] insects, nevertheless seems to have the most similarity with these insects. I cannot believe that this parasite, like that so-called Pediculus apis, should be a larva, although I am not completely certain given the infinite diversity of the larval forms. The hardness of the carapace, the perfectly formed walking legs, a certain, easily noticeable similarity in habitus with the Hippoboscids [keds], and in general a certain agreement with the bona fide dipterans, all speak for an adult developed form, in which I only observed this insect. Its relationship with the Diptera seems to be already evident from the mouth parts. I found at the mouth 1) two elongated, towards their end somewhat broadening, bristly organs, which I must hold for palps (actually maxillary palps, as they are those of all dipterans) and 2) between these palps an elongated, anteriorly divided into two narrow lobes, somewhat downward curved and extensible lower lip (a form of the so-called proboscis, as it shows for example in some Tipulis L.). By the way, this relationship is confirmed by the formation of the five tarsal leg segments, especially the adhesive lobes [aroliums] on them; by the almost spherical shape of the abdomen and even by the short, spine-like bristles on the whole body. The atrophy of the antennae and the absence of eyes, also occur at least in the Pupiparia.–However, the traits that differ from those of all other dipterans and would, consequently, be the main diagnostics of the group described here, which distinguish this parasite from these insects, if it is really one of them, are the following Braula traits: Four rudimentary antennae, namely two on each side [of the head] in place of the eyes, which are absent; a thorax divided into two segments similar to those of the abdomen; and instead of the pair of leg claws, a transverse row of numerous combs at the end of the last leg segment. [A rudimentary eye is likely confounded with an antenna].

9) I. genus. Braula, comb foot [common name]

3. Translation from Latin to English of the description of the species *Braula coeca* by Nitzsch (1818, p. 314-316)^2^

9) The genus I. Braula. ^[1]^) N.

Head tilted up and down, or inclined forward, triangular, flattened, with the mouth extending anteriorly and below.

Upper edge of the mouth (clypeus?) short, rounded at the front.

Lower edge somewhat curved, sloping, used for catching, bilobate, with rather narrow, elongated lobes [labella].

Palpi very small, thin and flat, elongated oval; with bristly edges.

Antennae: on each side [of the head] there is a pair of small, closely spaced projections, set in a furrow and covered with hairs; the projection set on the outside is larger, approximately at an angle, and cone-shaped; it is covered with an elongated, pointed, awl-shaped bristle, hairy, pinnate; the other, the one set on the inside, is smaller and is covered with a simple bristle. (So that is four rudimentary antennae). [A rudimentary eye is likely confounded with an antenna].

No eyes or ocelli.

Thorax, wingless, divided into two parts, short and as wide as the head; each of its two segments is similar to the segments of the abdomen.

Abdomen sessile, in line with the thorax, but widening, oval or rounded, convex, formed of four segments separated by thin sutures, somewhat mobile.

Tarsi perfect, bipedal [with two unguis (claws)?], hidden towards the thorax, with five articles; the last article, wider, comb-shaped underneath due to a transverse row of spines; with two oblong hairy terminal arolii.

Habitat[:] parasite on the honey bee.

Metamorphosis unknown.

Single species, which I recognized and called Braula coeca ("blind louse").

Observations: Extremely peculiar insect, quite different from the bee louse referred to by the authors (which is a beetle larva), about the size of a flea, but shaped like a hippoboscid fly [ked], or in some ways quite comparable to a small spider; the outer skin [lorica: literally "the armour"] is very hard, brown, shiny, yet almost everywhere made rough by short, sparse, stinger-like bristles. - Any one of the five bees captured in May and June and handed over to me by Cl. Keserstrinius, was feeding a Braula, which clung quite firmly to the thorax with its legs, so that it was difficult to unhook; the Braula was mostly motionless, but sometimes straightened the front part of its body and waved its front legs in a very peculiar way (which I also observed in Nycteribiidae). A Braula removed from its bee and placed on a sheet of paper or a glass slide would deftly run back and forth, anxiously searching for the bee's body, on which it would climb as soon as any part of the body had touched it, placing itself again in the same spot it had previously occupied. But when it had been completely separated from the bee for almost a quarter of an hour, its course was interrupted and soon, seized by a spasm, it violently contracted its legs; their movement continued weakening for about two hours and in the end it died. - Our Braula is probably not a larva. It is foreign to all orders of insects; however, I think it would be more relevant to add it to the order Diptera than to any other. But if it is not really related either to these Diptera, or to the Hymenoptera, to which it seems to be somewhat related, it is necessary for it to constitute a particular order of insects.

Note

Old French and German texts were initially translated with DeepL before being revised by Héloïse Bastide and Michael Lang, respectively. Latin description was translated to French by Christian Cheminade and then to English by DeepL and revised by Héloïse Bastide and Amir Yassin.

**References**

1.	de Réaumur, R.-A.F. (1740). Mémoires pour servir à l’histoire des insectes... Vol. 5 (de l’Imprimerie royale).

2.	Nitzsch, C. (1818). Die Familien und Gattungen der Thierinsekten (insecta epizoica); als Prodromus einer Naturgeschichte derselben. In.

1. ) βραύλα according to Hesychius of Alexandria means the same as φθείρ. [*phtheír* or louse in English as in Phthiraptera]
