## Supplementary material for "The genome of the blind bee louse fly reveals deep convergences with its social host and illuminates Drosophila origins": Document S2

**Supplementary text 2 – Analysis of potential horizontal transfer of transposable elements between the bee louse and the honey bee**

To detect possible HTT between *Braula coeca* and *Apis mellifera*, we used the *B. coeca* whole genome as query to perform a blastn similarity search against the whole *A. mellifera* genome (all default options, including “-task megablast”). All *B. coeca* genome regions longer than 299 bp and aligning to *A. mellifera* with an e-value lower than 0.0001 were extracted and clustered at 80% nucleotide identity threshold with vsearch^9^. The consensus sequence of each of the 50 resulting clusters were used as queries to perform blastx searches on the non-redundant protein database of NCBI using diamond^10^.

A total of eight consensus sequences had best hits to the *Famar1* element previously described in the earwig *Forficula auricularia*, known to be also present in *A. mellifera* as a result of horizontal transfer^11,12^. To verify that the *Famar1*-like element from *B. coeca* has indeed been involved in HTT, we compared the *Famar1-*like synonymous distance (dS) to a distribution of dS expected under vertical transmission since the last common ancestor of *B. coeca* and *A. mellifera* following the approach developed in ^13^.

This approach assumes that in case of HTT, TE dS should be much lower than dS expected under vertical transmission. Briefly, we calculated the dS over the transposase open reading frame between one copy of the *Famar1*-like element extracted from *C. coeca* and another copy of this element from the *A. mellifera* genome. We then compared this distance to the distribution of dS calculated over 2,179 alignments between genes or gene fragments that are deemed orthologous between the two species, *i.e.* single copy BUSCO^14^ genes that produce best reciprocal hits in blastp similarity searches^13^. We found that the *Famar1*-like dS (=0.12) fall below the 0.5% quantile (=1.76) of the distribution of dS calculated for orthologous genes (Supplementary Figure 2), confirming that the element has been acquired through horizontal transfer in *B. coeca* and *A. mellifera*. To assess whether the tight ecological interactions existing between *B. coeca* and *A. mellifera* might have favor direct transfer of this element between the two species, we assessed how closely related are *B. coeca* *Famar1*-like copies to those from *A. mellifera*. We first screened for the presence of this element in other animal genomes. We used the *Famar1* sequence^11^ as query to perform online blastn similarity searches (all default options, including “-task megablast”) on a total of 8180 animal genomes belonging to 11 insect orders as well as to Annelida, Chelicerata, Chiroptera, Cnidaria, Myriapoda, Nematoda, Platyhelminthes and Teleostei. We found full length copies showing >79% of nucleotide identity to this element in a total of 37 species. We aligned the ten copies from each genome the most similar to *Famar1* (or less when lower than ten copies were found) using Muscle^15^. The complete sequences of all *Famar1*-like copies included in this alignment are provided in Data SXXX. We then reconstructed a maximum-likelihood phylogeny of these copies using IqTree after nucleotide model detection using ModelFinder. Node support was quantified using ultrafast bootstrap (XXX) as implemented in IqTree.
