## Supplementary figures and images for "The genome of the blind bee louse fly reveals deep convergences with its social host and illuminates Drosophila origins"

### Figure S1

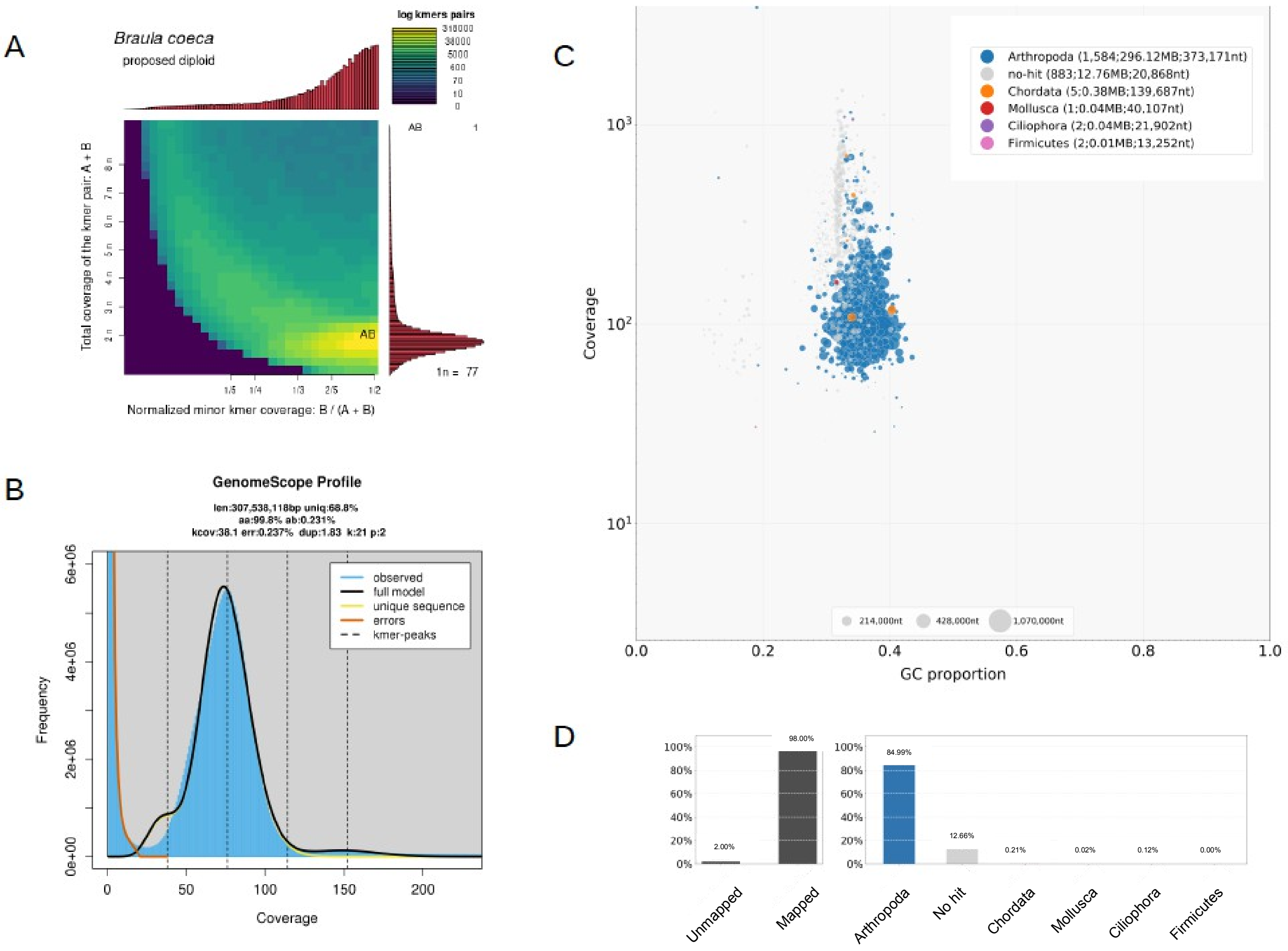

### Figure S2

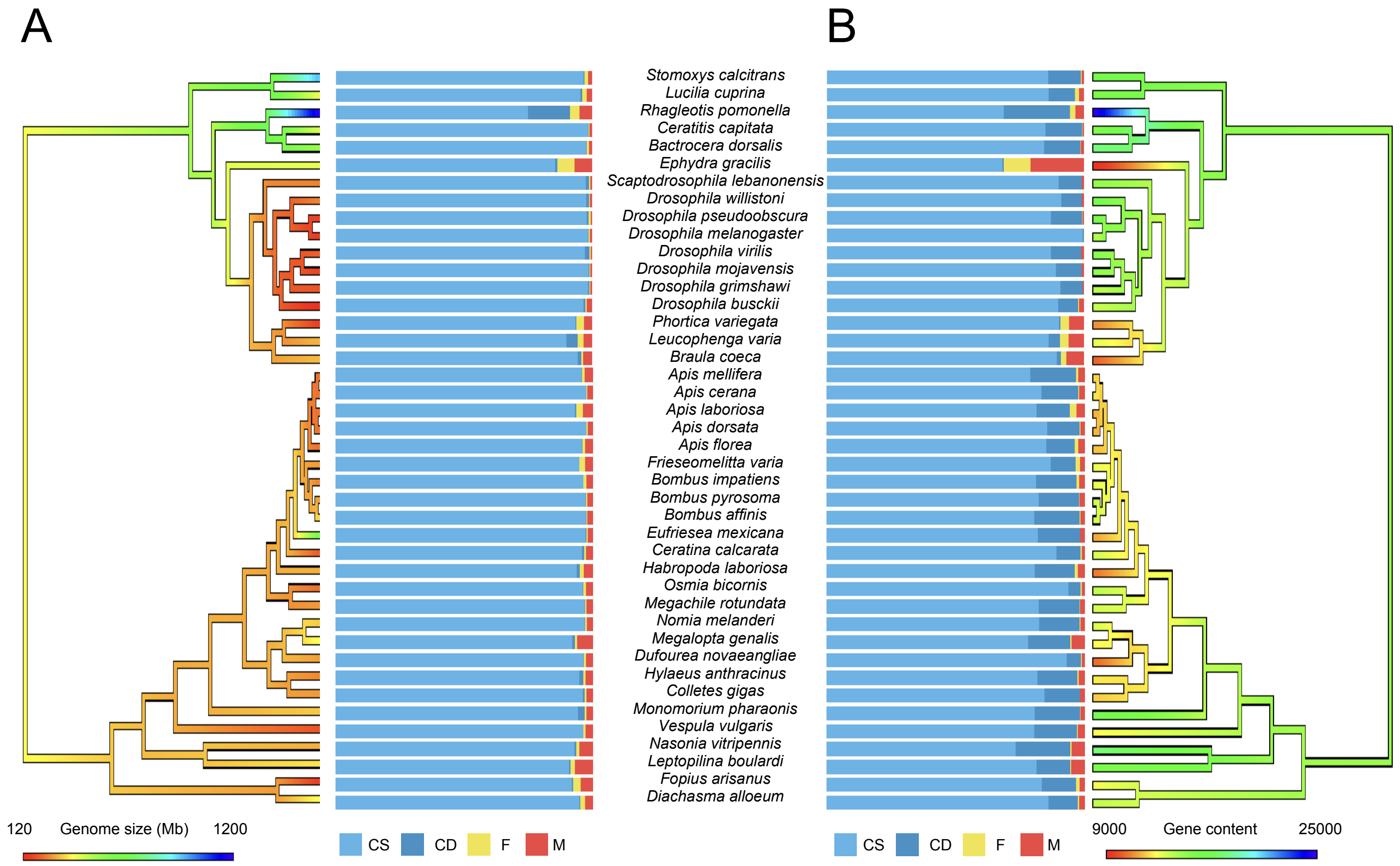

### Figure S3

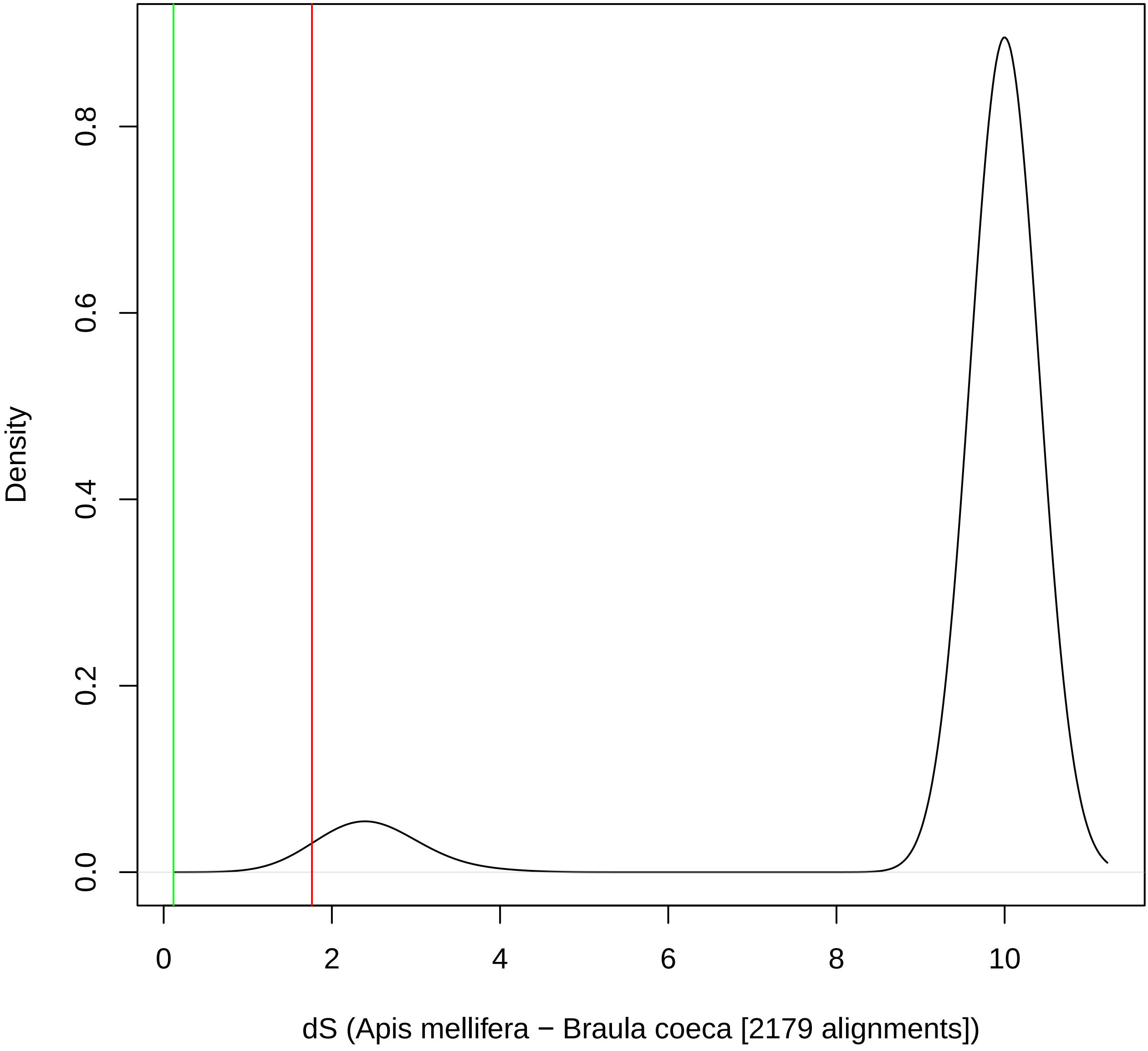

### Figure S4

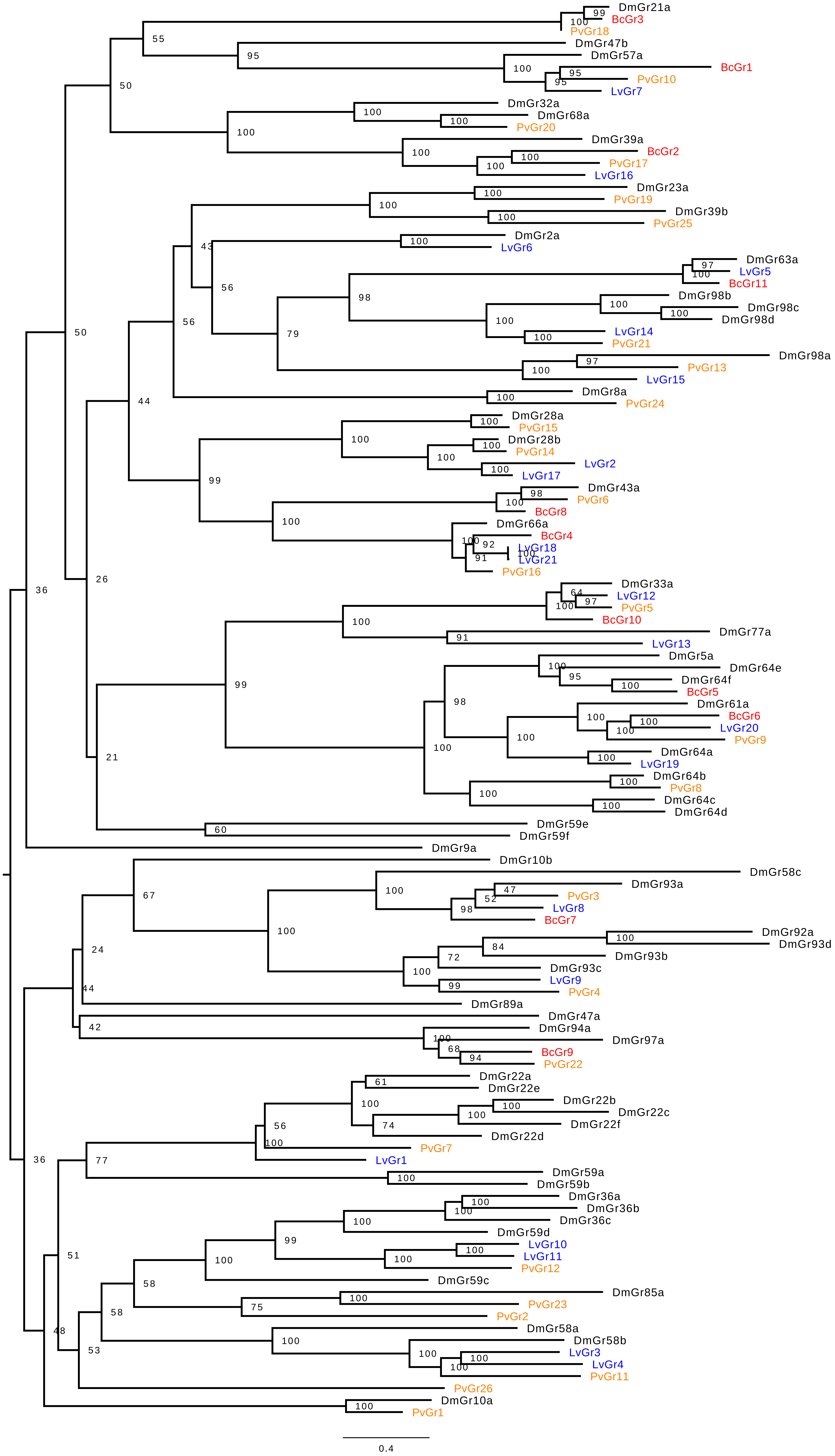

### Figure S5

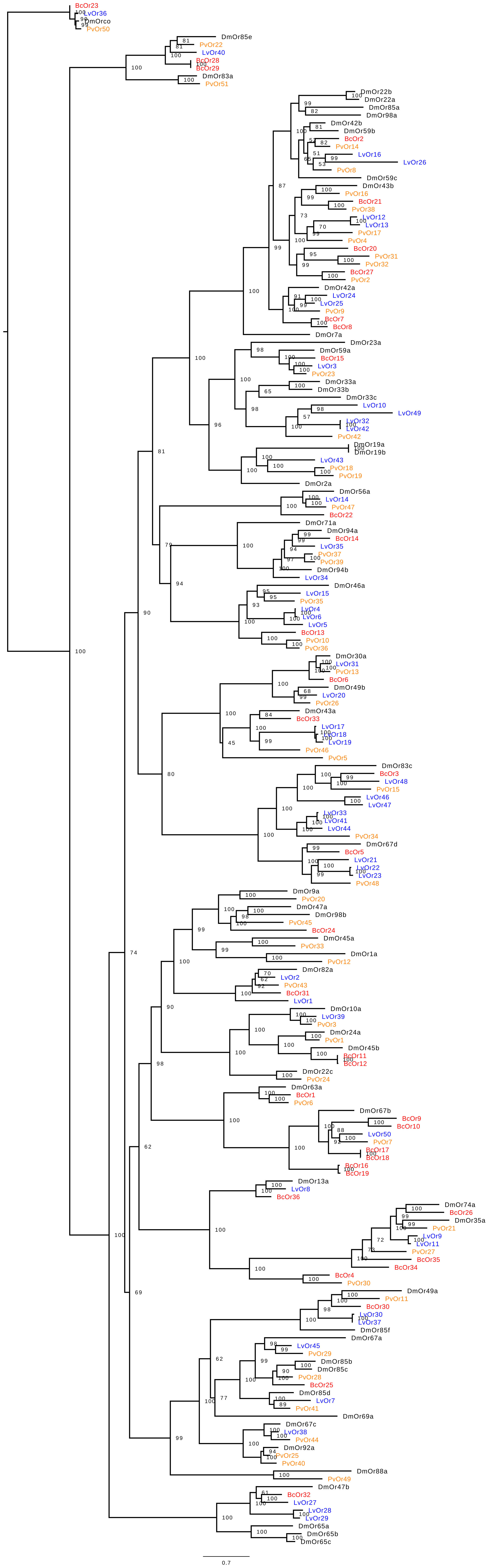
